# Mitochondrial dysfunction in Machado Joseph Disease: insights from a multi-model system

**DOI:** 10.1101/2025.05.01.651800

**Authors:** Ignacio Simo, Katherine J. Robinson, Julia Y. Kam, Andrea Kuriakose, Esmeralda Paric, Flora Cheng, Anastasiya Potapenko, Yousun An, Albert Lee, Angela S. Laird

## Abstract

Spinocerebellar ataxia type-3 (SCA3), also known as Machado Joseph disease, MJD) is a fatal, neurodegenerative disease belonging to the polyglutamine repeat disease family, caused by inheritance of an abnormal form of the ATXN3 gene, carrying a longer than usual trinucleotide repeat sequence. Within this study we explored mitochondrial function in a range of different experimental models of MJD, including transgenic zebrafish, mice and neuronal cells, to gain an understanding of mitochondrial function in the disease, and possible mechanisms of any dysfunction. Firstly, we examined the transgenic CMVMJD135 mouse model that develops impaired movement, neurodegeneration and decreased survival. We performed proteomic analysis on brain lysates extracted from a cohort of male and female WT and MJD mice, for analysis of differences in male and female mice separately. We identified that a major difference predicted by Ingenuity Pathway Analysis to be in both male and female MJD mice was related to impaired oxidative phosphorylation and mitochondrial dysfunction. We examined primary neuron cultures obtained from CMVMJD135 mice, validating the findings of the proteomic analysis, and finding changes to mitochondrial morphology, as well. We further examined a transgenic zebrafish model of MJD that expresses EGFP fused human ataxin-3 with short or long polyQ stretches (23 or 84Q, respectively) in neurons (expression driven under the elav/HuC promoter). The MJD zebrafish also exhibit altered mitochondrial electron transport chain complex protein levels, together with enhanced sensitivity to rotenone administration, which may be a valuable readout for treatment investigation studies in the future. Together, this study confirms, and extends on, the growing body of evidence suggesting that mitochondrial dysfunction plays a role in MJD, warranting investigation for therapeutic intervention.

## Introduction

Spinocerebellar ataxia type 3 (SCA3), also known as Machado-Joseph disease (MJD), is a progressive autosomal dominant neurodegenerative disorder caused by an expanded CAG trinucleotide repeat in the *ATXN3* gene. This results in a long polyglutamine (polyQ) tract within the ataxin-3 protein, leading to protein misfolding, aggregation, and cellular toxicity. MJD is characterised by symptoms of ataxia (loss of balance and coordination, resulting in an unsteady gait, clumsiness, and difficulties with fine motor tasks (1). Despite extensive research, the exact mechanisms driving neurodegeneration in MJD remain incompletely understood. Emerging evidence points to mitochondrial dysfunction as a central contributor to disease pathology, particularly in neurons, which are highly reliant on mitochondrial integrity for survival and function (2).

Mitochondria are critical for maintaining neuronal homeostasis, serving as the primary source of ATP production through oxidative phosphorylation, regulators of calcium dynamics, and mediators of apoptosis. Dysfunction in any of these processes can severely affect neuronal health, given the high metabolic demands of neurons and their limited capacity for regeneration. Mitochondrial deficits have been found to play a role in the pathogenesis and progression of a range of different neurodegenerative diseases, including Alzheimer’s disease (3, 4), Huntington’s disease (5) and amyotrophic lateral sclerosis (6). Mutations in genes such as superoxide dismutase 1 (SOD1), C9orf72, and CHCHD10 have not only been identified as causes of familial ALS, but have also been closely associated with mitochondrial dysfunction, suggesting that impaired mitochondrial homeostasis may play a central role in disease pathogenesis (7–9). Blockage of the mitochondrial electron transport chain, through administration of rotenone, also induces symptoms of Parkinson’s disease, including selective nigrostriatal dopaminergic degeneration and α-synuclein-positive cytoplasmic inclusions (10, 11). Moreover, striatal neurodegeneration linked to mitochondrial deregulation has been demonstrated in genetic and toxin-induced animal and cellular models and post-mortem HD human brain tissue (12, 13).

Recently, Harmuth *et al* (2022) reported that fibroblasts cultured from MJD patients exhibit abnormal mitochondria, resulting in decreased ATP (cellular energy) production (14). Hsu JY et alhave previously reported that truncated C-terminal Fragments of Mutant ATXN3 disrupts the mitochondrial electron transport chain (ETC), leading to impaired oxidative phosphorylation and ATP production (15). Proteomic analyses of brain tissues from MJD models have previously revealed downregulation of several ETC complexes, including complex I and IV, further implicating energy metabolism defects in the disease process (check ref).

Mitochondrial dysfunction in MJD is not limited to impaired energy production. Morphological abnormalities, such as mitochondrial fissions, suggests that mitochondrial dynamics disruption is a common characteristic in models expressing a truncated ATXN3 form which generate aggregates (15). These structural defects are accompanied by reduced levels of key mitochondrial proteins such as Mfn-1 and Mfn-2Morphological abnormalities. Furthermore, mitochondrial trafficking, a process critical for distributing mitochondria to regions of high metabolic demand in neurons, appears to be compromised in MJD. Impaired trfficking disrupts synaptic function and likely contributes to the progressive degeneration observed in MJD (16).

In this study, we investigate the mitochondrial dysfunction associated with MJD using a combination of in vivo and in vitro models, including zebrafish, primary mouse neuronal cultures, and brain tissues from MJD mice. Using proteomic analyses, mitochondrial staining, and functional assays, we characterize the relationship between polyQ expansion length and mitochondrial impairment. We hypothesize that longer polyQ repeat lengths exacerbate mitochondrial dysfunction, as evidenced by altered morphology, reduced protein levels, and increased sensitivity to mitochondrial stressors. By elucidating these mechanisms, this work aims to enhance our understanding of mitochondrial contributions to MJD pathogenesis and provide a foundation for therapeutic interventions targeting mitochondrial dysfunction in polyQ diseases.

Mitochondria are critical for maintaining neuronal homeostasis, serving as the primary source of ATP production through oxidative phosphorylation, regulators of calcium dynamics, and mediators of apoptosis. Dysfunction in any of these processes can severely affect neuronal health, given the high metabolic demands of neurons and their limited capacity for regeneration. In neurodegenerative diseases such as MJD, disruptions in mitochondrial dynamics, including fusion, fission, trafficking, and quality control, are consistently observed (17, 18)

Furthermore, mitochondrial trafficking, a process critical for distributing mitochondria to regions of high metabolic demand in neurons, appears to be compromised in MJD. Impaired trafficking disrupts synaptic function and contributes to the progressive degeneration observed in disease models These findings align with observations from other polyQ diseases, such as Huntington’s disease, where similar disruptions in mitochondrial transport and dynamics are reported (check).

In this study, we investigate the mitochondrial dysfunction associated with MJD using a combination of in vivo and in vitro models, including zebrafish, primary mouse neuronal cultures, and brain tissues from MJD mice. Using proteomic analyses, mitochondrial staining, and functional assays, we characterize the relationship between polyQ expansion length and mitochondrial impairment. We hypothesize that longer expansions exacerbate mitochondrial dysfunction, as evidenced by altered morphology, reduced protein levels, and increased sensitivity to mitochondrial stressors. By elucidating these mechanisms, this work aims to enhance our understanding of mitochondrial contributions to MJD pathogenesis and provide a foundation for therapeutic interventions targeting mitochondrial dysfunction in polyQ diseases.

## Materials and Methods

### Animal maintenance

All animal experiments were conducted in accordance with the Australian Code for the Care and Use of Animals for Scientific purposes (8^th^ Edition, 2013) and approved by the animal ethics committee of Macquarie University (ARAs: 2016/004, 2017/044, 2017/003and 2017/019).

Zebrafish were maintained under standard conditions in a recirculating aquarium system at 28.5°C with a 13-h light / 11-h dark cycle. Zebrafish embryos used in this study were obtained from transgenic MJD zebrafish generated as previously described (19). This line was established by crossing the Tg(elav3:Gal4-VP16; mCherry) driver line with responder lines Tg(UAS:dsRED,EGFP-ATXN3_Q23) or Tg(UAS:dsRED,EGFP-ATXN3_Q84), allowing neuronal expression of human ataxin-3 (hATXN323Q or hATXN384Q) fused to EGFP. F1 zebrafish were used for further analysis.

Transgenic CMVMJD135 mice expressing mutant human *ATXN3* under a CMV promoter were bred and housed at Australian BioResources (Moss Vale, Australia) under pathogen-free conditions. Transgenic mice expressing human mutant *ATXN3* (CMVMJD135 mice) were bred with wildtype C57Bl6 mice to obtain heterozygous *ATXN3* expressing mice and non-transgenic littermate controls. Transgene expression was confirmed via PCR and CAG repeat expansion length was confirmed via fragment analysis.

This study utilises a cohort of mice previously described by Gamage et al. (20). Briefly, a total of 55 male and female mice were delivered to the Central Animal Facility at Macquarie University and allowed to acclimate for 7 days prior to commencing experimentation. Animals were group-housed according to sex and genotype in exhaust ventilated cages with ad libitum access to standard chow and water. Mice were provided red domes or cardboard tunnels for environmental enrichment, and housing rooms were maintained at 21°C with a 12-hour light/dark cycle. All animals under weekly monitoring to assess wellbeing and neurological symptoms. The presence or absence of neurological symptoms such as hindlimb reflex, tremor and gait abnormalities were scored on a 4-point scale, with a score of 0 indication no neurological symptoms and a score of 4 indicating severe neurological symptoms. CMVMJD135 mice were deemed to reach humane endpoints upon reaching a total neurological score of 10 and were euthanised (age range: 20-25 weeks of age). Non-transgenic littermates were euthanised at 25 weeks of age.

### Drug treatment

hATXN3 23Q and 84Q zebrafish embryos were obtained from three independent matings and, at 6 hours post-fertilization (hpf), randomly divided into three treatment groups: DMSO, 100 μM NaN_3_, or 200 nM rotenone (Sigma; prepared in DMSO). Treatments were carried out continuously for 3 days. During this period, embryos were imaged daily, and their developmental stages were assessed and compared across treatment conditions.

### Protein Extraction from Cerebellum

Mice were deeply anaesthetised via injection of sodium pentabarbitone (100mg/kg, i.p.) and subjected to intracardiac perfusion with ice-cold phosphate buffer saline (PBS). Brains were extracted and a sagittal hemi-section was performed. One hemisphere of brain was immediately snap frozen in liquid nitrogen and stored at −80°C until processing.

The cerebellum was dissected and placed in RIPA buffer (5μL/mg) containing cOmplete Protease Inhibitor Cocktail (Roche) and PhosSTOP Phosphatase Inhibitor Cocktail tablets (Roche). Cerebellum tissue was homogenised via sonication using an Omniruptor 250 Ultrasonic Homogeniser (Omni International). Protein lysates were centrifuged at 13200 x *g* for 20 minutes at 4°C. After centrifugation, the clear supernatant was collected and used for further analysis. Protein concentration was determined using a Pierce BCA Protein Assay Kit (Thermo Fisher Scientific).

### In-gel digestion

*2*0 μg of brain protein lysates were digested following the protocol (21). In short, the lysates were boiled at 95°C for 5 minutes in 1x loading buffer (1610747, Biorad) containing 1x reducing reagent (NP0004, Invitrogen). The mixture was run 3 cm into a 4-12% SDS-PAGE gel (WG1402BOX, Invitrogen) and excised from the gel into 3 fractions. Proteins were reduced with 10 mM dithiothreitol (D0632, Sigma), alkylated with 55 mM iodoacetamide (I1149, Sigma) then digestion with trypsin (1:50, enzyme:protein, 90057, Thermo Fisher Scientific) overnight at 37°C. Formic acid (5330020050, Sigma) was added to a final concentration of 1% to inactivate the digestion. The extracted tryptic peptides were desalted on a pre-equilibrated C18 Omix tip (A57003100K, Agilent) and eluted using 50% (v/v) acetonitrile (1000292500, Sigma) in 0.1% formic acid. Lyophilised peptides were resuspended in 0.1% formic acid for liquid chromatography-tandem mass spectrometry (LC-MS/MS) analysis.

### Reverse phase liquid chromatography tandem mass spectrometry

The peptides were separated on a Ultimate 3000 nanoLC (Thermo Fisher Scientific) fitted with the Acclaim PepMap RSLC column (2 μm particle size, 0.075 mm diameter and 150 mm length, 164534, Thermo Fisher Scientific), using a 60 mins gradient (2–80% v/v acetonitrile, 0.1% v/v formic acid) running at a flow rate of 300 nl/min. The electrospray source was fitted with a 10 μm emitter tip and maitenaned at 1.6 kV voltage. Peptides eluted from the nano LC column were ionised into the Q-Exactive Plus mass spectrometer (Thermo Fisher Scientific). Precursor ions were selected for MS/MS fragmentation using a data-dependent “Top 10” method operating in Fourier Transform (FT) acquisition mode with higher-energy C-trap dissociation (HCD) fragmentation. FT-MS analysis was carried out at 70,000 resolution and with an automated gain control (AGC) target of 1 × 10^6^ in full MS. MS/MS scans were carried out at 17,500 resolution with an AGC target of 2 × 10^4^ ions. Maximum injection times for full MS and MS/MS are set at 30 and 50 milliseconds, respectively. The charge exclusion was set to unassigned and 1+ charged state with a dynamic exclusion of 20 seconds. The ion selection threshold for triggering MS/MS was set to 25,000 counts, and an isolation width of 2.0 Da was used to perform HCD fragmentation with a normalised collision energy of 27.

### Data analysis

Raw spectra files were processed using Proteomie Discoverer software 2.4 (Thermo Fisher Scientific) incorporating the SEQUEST search algorithm. Peptide identification was determined using a 20 ppm precursor ion tolerance and a 0.1 Da MS/MS fragment ion tolerance for FT-MS and HCD fragmentation. Carbomidomethylation static modification (+57.021 Da) of cysteines and oxidation (+15.995 Da) of methionine, N-terminal acetyl (+42.011 Da) variable modifications were set with a maximum of two missed cleavages allowed. The data were processed using Percolator to estimate false discovery rates (FDR), with protein identifications validated at a q-value threshold of 0.01. Label-free quantification (LFQ) was performed using intensity-based methods in Proteome Discoverer 2.4, applying the software’s default parameters. Briefly, peptide spectral matches (PSMs) were filtered using a maximum delta Cn of 0.05, a rank of 0, and a delta mass of 0 ppm. PSM and peptide validation was carried out using a strict FDR or 0.01, and a relaxed FDR of 0.05. Peptides shorter than 6 amino acids were excluded from the analysis.

For each sample set, chromatographic alignment of PSMs was conducted with a mass tolerance of 10 ppm and a maximum retention time shift of 10 minutes. Peptide groups used in protein quantification followed default settings, where a peptide was considered unique if present in only one protein group. Quantification was based on both unique and razor peptides (those shared among multiple protein groups), using precursor ion intensity.

Protein abundance was determined by summing the abundances of individual peptide groups, and protein ratios were calculated using pairwise comparisons based on the geometric median of peptide group ratios. Statistical significance was assessed using Student’s t-test, comparing each peptide or protein against the background distribution of all ratio values. P-value were adjusted for multiple testing using Benjamini-Hochberg correction.

### Primary neurons culture

Pregnant female mice were euthanised via cervical dislocation on gestational day 16. Primary neuronal cultures were prepared as per Fath et al 2009 (22). Embryos were extracted and anaesthetised in ice cold Hanks buffered Saline solution. The cortex and hippocampi were dissected from the brains of mixed gender and genotype embryonic mice at day 16 of gestation (E16) and dissociated via mechanical trituration following exposure to trypsin and DNase I. Neurons were plated at a density of X cells in PDL-coated well plates containing Dulbecco’s Modified Eagle Medium (DMEM) with 10% fetal bovine serum. At two-hours post-plating, DMEM was aspirated and replaced with neurobasal media containing 2% B27 supplements and 2mM Glutamax. Neurons were maintained in a 37°C incubator with 5% CO_2_.

### Western blot

Primary cells were cultured in a 6-well plate with 300,000 cells per well at 18 days in vitro (DIV). The cells were lysed with 30 µL of RIPA buffer containing cOmplete Protease Inhibitor Cocktail (Roche) and PhosSTOP Phosphatase Inhibitor Cocktail tablets (Roche) per well. After scraping the cells, the lysates were transferred to a new well, scraped again (having a final number of cells of 600000 cells per tube). The supernatant containing the RIPA-soluble proteins was collected for analysis. For zebrafish samples, 20 zebrafish (5 days post-fertilization, dpf) were used per replicate. The yolk was removed following the procedure described in (23). **T**he remaining tissue was homogenized in RIPA buffer containing cOmplete Protease Inhibitor Cocktail (Roche) and PhosSTOP Phosphatase Inhibitor Cocktail tablets (Roche). In both cases, the homogenized samples were then centrifuged at 13,000 rpm for 15 minutes at 4°C, and the supernatant containing the RIPA-soluble proteins was collected for analysis. The total protein concentration was determined using a Pierce BCA Protein Assay Kit (ThermoFisher Scientific) according to the manufacturer’s instructions. For Western blotting, 20 µg of protein from each sample was prepared with NuPage Sample Reducing Agent (Invitrogen) containing dithiothreitol and Laemmli Sample buffer (Bio-Rad), denatured, and separated using a 4-12% acrylamide gradient NuPAGE Bis-Tris gel (Invitrogen). Proteins from both primary cell and zebrafish samples were transferred to a 0.45 µm polyvinylidene difluoride (PVDF) membrane (Cytiva) and incubated overnight in TBS-T 0.1% BSA 5% with specific primary antibodies: anti-mtND5 (1:2000; Abcam ab15895) (Abcam, check catalogue), anti-SDHA (1:3000; Cell Signalling, D6J9M), anti-VDAC1 (1:2000), anti-Pyruvate Kinase (1:2000; Abcam ab156849). The immunoblots were incubated with the corresponding secondary antibodies: goat anti-rabbit IgG (H+L) HRP Conjugate (Promega, #W4011) for rabbit primary antibodies, and goat anti-mouse IgG (H+L) HRP Conjugate (Promega, #W4021) for mouse primary antibodies. The protein bands were visualized using the ImageQuant LAS 4000 (Cytiva) by chemiluminescent reaction with Clarity Western ECL Substrate (Bio-Rad). The images were exported as digital 8-bit TIFF files and analysed using ImageStudio (LI-COR Biosciences). Quantification of the target proteins was performed, with normalization to GAPDH (for zebrafish samples) or β-actin (for primary cell samples), which were used as loading controls

### Succinate assay

Succinate concentrations were assessed by colorimetry using commercially available kits (ab204718, Abcam). Transgenic zebrafish expressing 23Q or 84Q human ataxin-3 were maintained at 28.5°C in standard E3 medium. At 5 days post-fertilization (dpf), pools of 50 embryos were prepared for each biological replicate, with 6 replicates analysed per genotype. Embryos were rinsed in extraction buffer and homogenized according to the instructions provided by the succinate assay kit manufacturer. Mice brain tissues were collected from wild-type (WT) and MJD mutant (CMVMJD135) mice. For each condition, three biological replicates were used, with each replicate consisting of 100 mg of brain tissue. Brain tissues were homogenized in the extraction buffer provided in the kit using a dounce homogenizer glass Potter-Elvehjem on ice and centrifuged at 10,000 × g for 10 minutes to remove debris. The supernatants were collected and stored at −80°C for subsequent analysis. Homogenized zebrafish and mouse brain samples were deproteinized using 10 kDa molecular weight cutoff spin filters (Abcam, ab93349) to eliminate interfering proteins. The filtrates were diluted in the assay buffer provided in the kit to ensure absorbance readings fell within the linear range of the standard curve. Succinate levels were measured by a colorimetric assay that detects succinate through a coupled enzymatic reaction producing a signal at 450 nm. A standard curve was prepared with succinate concentrations ranging from 0 to 10 nmol/well. Fifty microliters of each sample or standard were added to a 96-well plate in triplicate, and the reaction mix was prepared and added to each well per the manufacturer’s instructions. Plates were incubated at 37°C for 30 minutes, protected from light, and the absorbance at 450 nm was measured using a CLARIOstar Plus Microplate Reader (BMG Labtech). Succinate concentrations were calculated from the standard curve using linear regression and normalized to protein content for zebrafish samples or tissue weight for mouse samples.

### DNA quantification

Total DNA, including mitochondrial DNA, was extracted from primary cells cultured in 6-well plates (350,000 cells per well) by adding 300 μL of TRIzol™ reagent (Invitrogen) directly to the wells. DNA was isolated following the manufacturer’s protocol, ensuring recovery of both nuclear and mitochondrial DNA for downstream analysis. Mitochondrial DNA (mtDNA) levels were quantified in primary neurons derived from WT and mutant MJD mice. Quantitative PCR (qPCR) was performed to determine mtDNA levels. Primers specific to mitochondrial DNA (target gene) and GAPDH (reference gene) were designed to ensure amplification efficiency. The reactions were carried out using SYBR Green PCR Master Mix (Applied Biosystems) in a StepOnePlus Real-Time PCR system (Applied Biosystems). Each reaction consisted of 10 µL of SYBR Green mix, 0.2 µM of each primer, 2 µL of DNA template, and nuclease-free water to a final volume of 20 µL. The thermocycling conditions included an initial denaturation step at 95°C for 5 minutes, followed by 40 cycles of 95°C for 15 seconds and 60°C for 1 minute. Relative mtDNA levels were calculated using the 2^-ΔΔCt method (reference), with GAPDH serving as the internal reference for normalization. All reactions were performed in triplicate, and no-template controls were included to confirm the absence of contamination.

### Mitochondrial labelling by Mitotracker

For mitochondrial staining, primary cells cultured on coverslips in 24-well plates at 18 days in vitro (DIV) were incubated with 500 µL of MitoTracker Red CMXRos at a final concentration of 100 nM in Neurobasal medium without serum for 30 min at 37°C. Following incubation, cells were washed twice with PBS and fixed for 15 min in 3.7% paraformaldehyde at room temperature. After two additional PBS washes, coverslips were mounted with Hoechst-containing mounting media for subsequent confocal microscopy analysis.

### Image deconvolution, Image processing and thresholding

For image deconvolution, processing, and thresholding, we utilized the Mitochondrial Analyzer plugin in ImageJ (24). Briefly, mitochondrial morphology was analysed from 2D images acquired using [microscope model] with a [63X objective lens details] using a zoom 3X at [resolution details]. Raw images were pre-processed to enhance mitochondrial signal and reduce background noise: Background subtraction (radius = 1 µm) to minimize non-mitochondrial fluorescence. Sigma filtering (radius = 0.1 µm, sigma = 2.0) to smooth signal while preserving edges. Contrast enhancement (block size = 64, slope = 2.0) to improve visualization of dim structures. Gamma correction (value = 0.80) to normalize intensity across images. Mitochondria were segmented using adaptive thresholding (block size = 1.25 µm, empirically determined C value). Post-processing steps included despeckling, removal of outliers (radius = 0.15 µm), and binary mask generation for final segmentation. Using the Mitochondrial Analyzer plugin, we extracted key parameters including Area and perimeter, Form factor (FF) and aspect ratio (AR) (to evaluate mitochondrial elongation and branching) and Branching metrics (to quantify mitochondrial network complexity).

### Statistics

Data analysis was performed using GraphPad Prism (version 10) software. Group comparisons were made using one-way ANOVA, followed by a Tukey’s post-hoc test to identify differences. In cases in which only two groups were compared, comparisons were made using Student’s t-test. All graphs display group mean data ± s.e.m.

## Results

### Proteomic analysis reveals differentially regulated pathways in brains of WT and MJD Mice

To analyse expression changes and altered metabolic pathways in the brains of mice expressing a mutant copy of 84Q hATXN3, we performed global proteomic analysis of male and female mutant and WT mice. Raw files were searched using Proteome Discoverer 2.4 (Thermo Fisher Scientific) against the Mus Musculus Uniprot database (Taxon ID 10090) and the protein list was filtered (q-value < 0.01, protein abundance cut-off of ≤ −2 or ≥ 2-fold with an adjusted p-value < 0.05) for Ingenuity Pathways analysis. Proteomic analysis revealed widespread dysregulation of several cellular pathways, with a notable impact on mitochondrial function (Figure 1). A significant proportion of the altered pathways were downregulated (XX%), while upregulated pathways were found to be (YY%), highlighting a complex and profound alteration in metabolic processes in MJD. Among these, mitochondrial dysfunction, particularly within the oxidative phosphorylation (OXPHOS) pathway, was most significantly affected.

**Figure 1:**
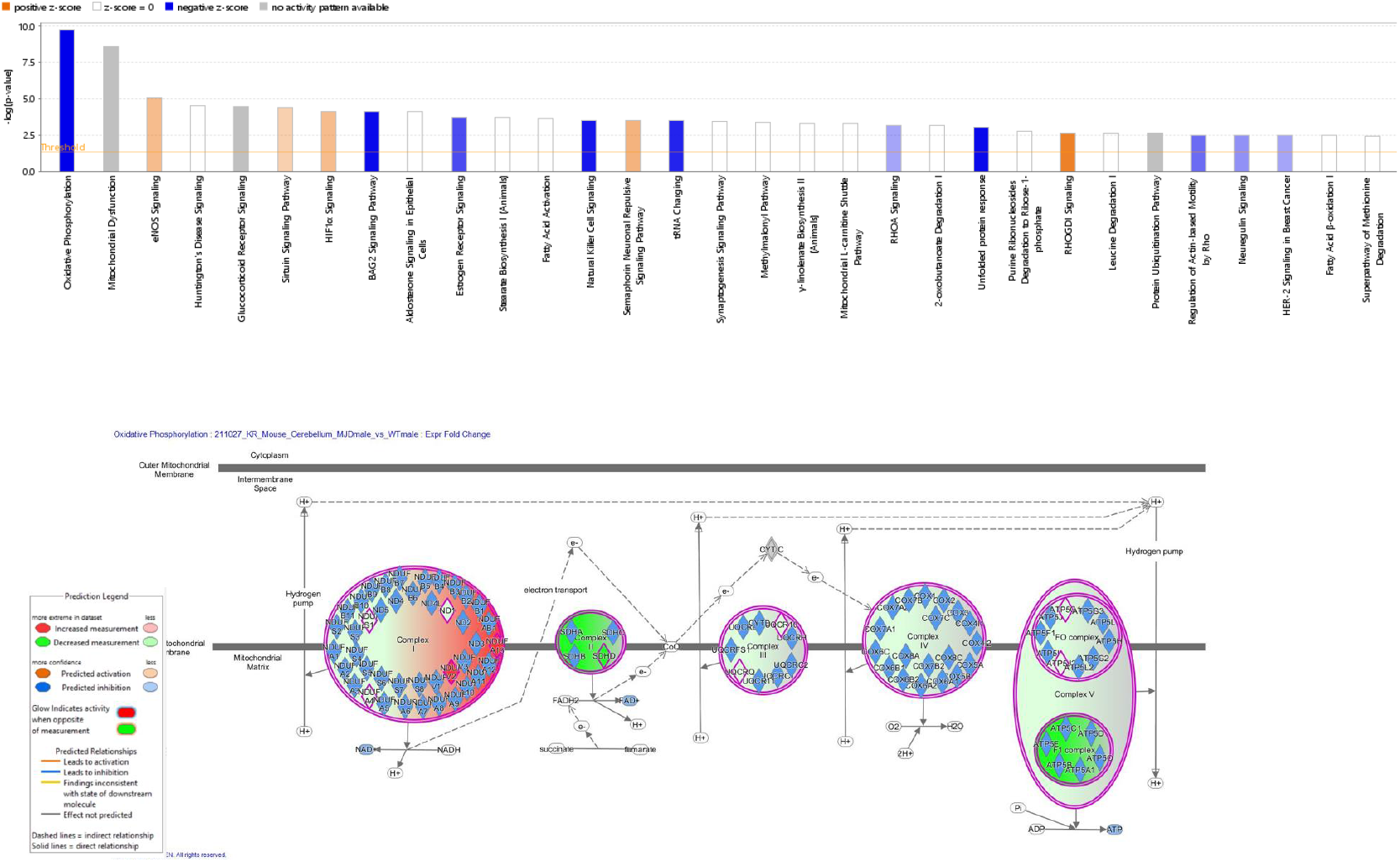

Within the OXPHOS pathway, the most pronounced reductions were observed in complexes II (succinate dehydrogenase) and V (ATP synthase). These complexes showed decreased expression of multiple subunits in MJD brains compared to WT controls. Notably, these results were obtained from male mice; however, parallel analysis in female cohorts revealed similar patterns of mitochondrial protein downregulation. This consistency across sexes suggests that mitochondrial and electron transport chain (ETC) dysfunction is a robust and sex-independent feature of the disease. The marked reduction of complex II, which links the tricarboxylic acid (TCA) cycle with the ETC, indicates disrupted electron flow and compromised energy production. Likewise, the downregulation of complex V components points to impaired ATP synthesis capacity. Together, these findings reinforce mitochondrial dysfunction, particularly at the level of OXPHOS, as a central and early hallmark of neurodegeneration in MJD.

### Western blot validation of proteomic findings in MJD neurons

To validate the proteomic results, we performed Western blot analysis on proteins extracted from primary neurons derived from WT and MJD mice after 18 days in vitro. Consistent with our proteomic data, we observed a significant reduction in SDHA (a subunit of Complex II) and mtND5 (a subunit of Complex I), confirming deficiencies in key components of the electron transport chain (Figure 2 B, C). Additionally, pyruvate dehydrogenase, a crucial enzyme linking glycolysis to mitochondrial respiration, was also decreased in MJD neurons, suggesting broader impairments in mitochondrial metabolism (Figure 2D).

**Figure 2:**
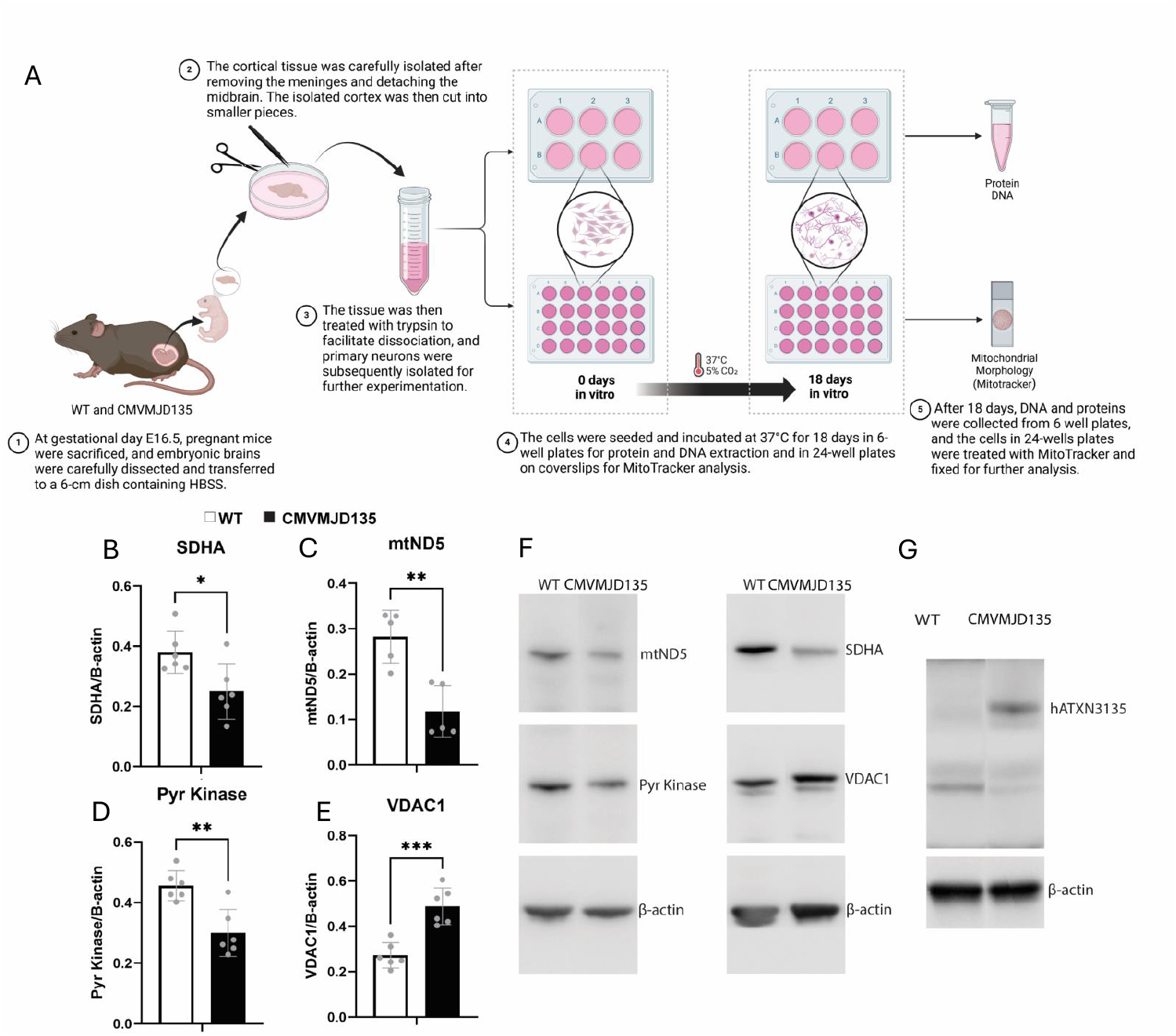
Primary cortical neuron cultures from MJD and WT embryos and analysis of mitochondrial and metabolic markers. (A) Schematic overview of the primary neuron culture process. Cortical neurons were isolated from embryonic brains of pregnant MJD and WT mice, cultured for 18 days in vitro (DIV), followed by protein or DNA extraction, and MitoTracker analysis. (B–E) Quantification of mitochondrial and metabolic markers by Western blot in primary cortical neurons: (B) SDHA, (C) mtND5 (n=5), (D) Pyruvate kinase (PK), and (E) VDAC1. (F) Representative Western blot images corresponding to the quantifications shown in (B–E). (G) Detection of mutant ATXN3 protein in primary neurons from MJD mice; ATXN3 was not detected in WT samples. β-actin was used as the loading control in all Western blots. Data were obtained from six biological replicates, except for mtND5 (n=5).

In contrast, VDAC1 was significantly upregulated in MJD neurons (Figure 2E). Given its role in regulating mitochondrial permeability and apoptosis, this increase may be associated with mitochondrial dysfunction and elevated oxidative stress.

### EMect of mutant ATXN3 on mitochondrial morphology and biomass

Fluorescence microscopy and quantitative image analysis revealed significant alterations in mitochondrial morphology in MJD primary neurons at 18 days in vitro (DIV) compared to WT (Figure 3A). Specifically, mitochondria in mutant neurons exhibited a significant reduction in overall size and structural complexity. We observed decreases in mitochondrial area, perimeter, form factor, longest shortest path, as well as multiple branching parameters, including the number of branches, total branch length, mean branch length, branch endpoints, and branch junctions (Figure 3B-J. These changes suggest that mitochondria in MJD neurons are more fragmented and less interconnected, indicating potential defects in the balance of mitochondrial fission and fusion, processes critical for maintaining mitochondrial network integrity and function.

**Figure 3.**
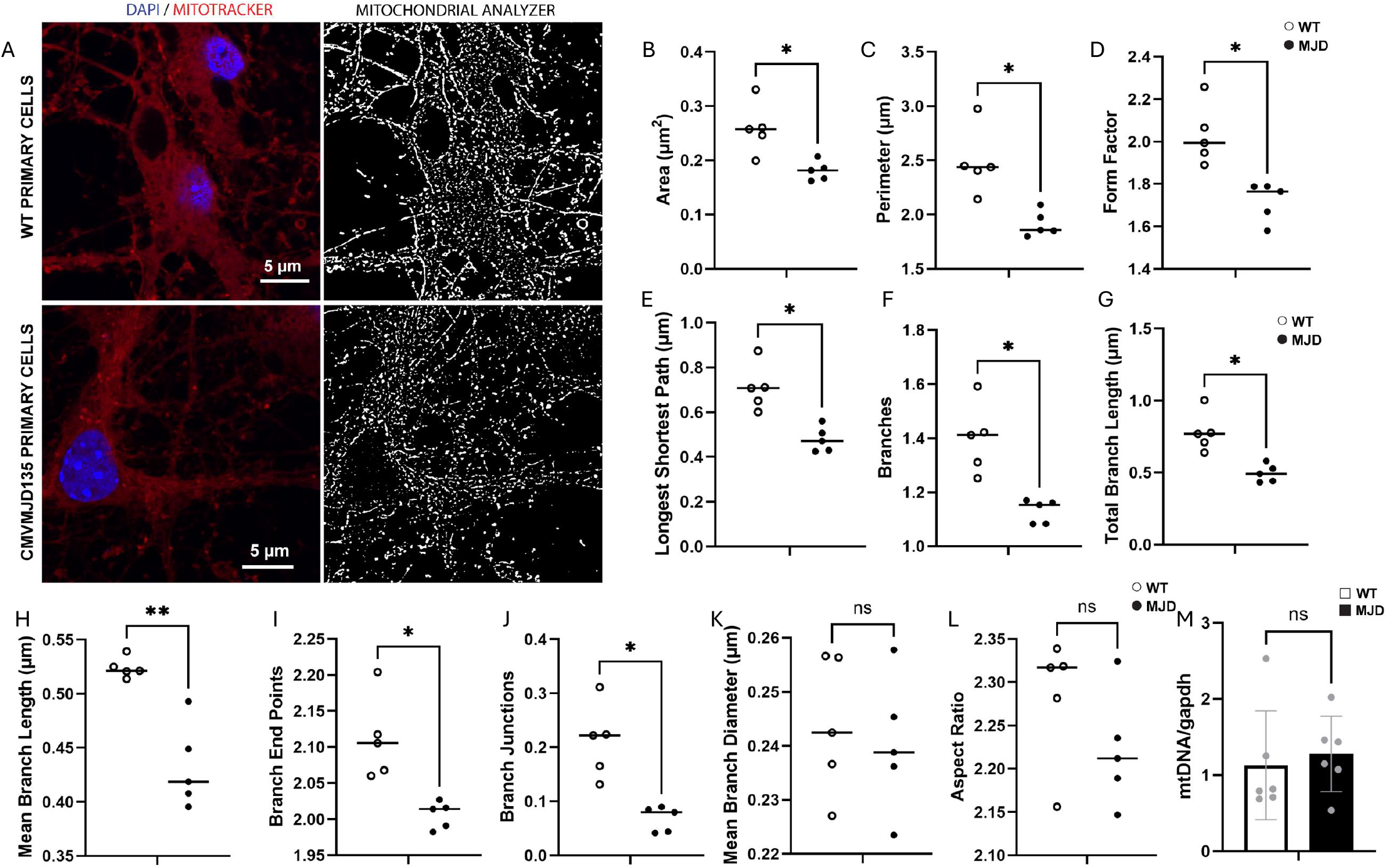
Analysis of mitochondrial morphology and mtDNA content in primary cortical neurons from WT and MJD mice. (A) Representative confocal images of primary cortical neurons stained with MitoTracker Red (mitochondria) and DAPI (nuclei). Images include processed outputs from the mitochondrial morphology analyzer in ImageJ. (B–L) Quantification of mitochondrial morphology parameters in WT and MJD neurons: (B) area, (C) perimeter, (D) form factor, (E) longest shortest path, (F) number of branches, (G) total branch length, (H) mean branch length, (I) number of branch endpoints, (J) number of branch junctions, (K) mean branch diameter, and (L) aspect ratio. (M) Mitochondrial DNA (mtDNA) quantification by qPCR. GAPDH was used as the nuclear reference gene. All analyses were performed using six biological replicates per group (WT and MJD). Asterisks indicate statistically significant differences (P < 0.05).

In contrast, no significant differences were observed in the mean branch diameter and aspect ratio or between MJD and WT neurons (Figure 3K and L), suggesting that while mitochondrial size and branching complexity are compromised, their overall shape remains largely unchanged.

To evaluate whether mitochondrial number was affected, mitochondrial DNA (mtDNA) levels were quantified by qPCR using GAPDH as a reference gene. No significant differences in mtDNA levels were observed between mutant and WT neurons, suggesting that the mitochondrial biomass remains unchanged in disease (Figure 3 M). This indicates that the observed changes in mitochondrial morphology are not due to alterations in mitochondrial quantity but are likely a reflection of compromised mitochondrial quality or function.

### Genotype-dependent sensitivity of ATXN3 23Q and ATXN3 84Q zebrafish embryos to electron transport chain disruptions: differential responses to Complex I and IV inhibition

To assess the developmental responses of ATXN3 23Q and ATXN3 84Q zebrafish embryos, we treated them with DMSO (control), sodium azide (a cytochrome c oxidase inhibitor targeting complex IV), and rotenone (a selective Complex I inhibitor). Sodium azide treatment resulted in comparable developmental delays in both ATXN3 23Q and ATXN3 84Q embryos, indicating that the inhibition of complex IV similarly affects embryonic development regardless of the genetic background (Figure 4A). This suggests that both genotypes exhibit an equal tolerance to disruptions in the terminal stage of the electron transport chain.

**Figure 4.**
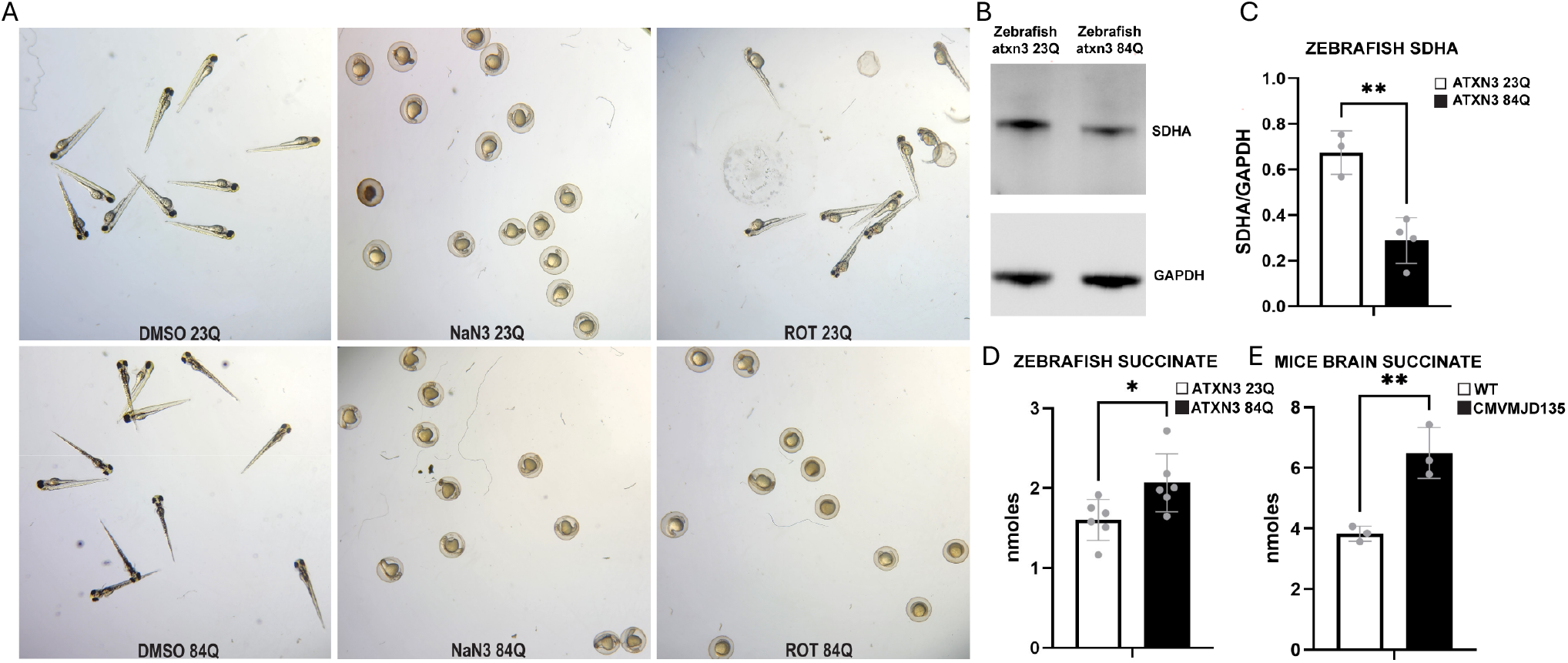
Impact of polyQ expanded ATXN3 on mitochondrial function and succinate accumulation in zebrafish and mouse brain. (A) Sensitivity of hATXN3 23Q and 84Q zebrafish embryos to mitochondrial inhibitors NaN_3_ and rotenone, compared to DMSO controls. (B) Western blot analysis of SDHA levels in hATXN3 23Q and 84Q zebrafish. GAPDH was used as the loading control (n = 3 for hATXN23Q embryos; n = 4 for hATXN384Q embryos). (C) Representative image of succinate assay in mouse brain tissue. (D–E) Quantification of succinate levels (nmol): (D) in hATXN3 23Q and 84Q zebrafish (n = 6 per group), and (E) in WT and MJD mouse brains (n = 3 per group). White bars represent WT samples and dark bars represent MJD model samples. Asterisks indicate statistically significant differences (P < 0.05).

In contrast, treatment with rotenone revealed genotype-dependent differences. While both groups did not experience developmental delays compared to DMSO-treated controls, ATXN3 84Q embryos showed a significantly more pronounced delay than ATXN3 23Q embryos (Figure 4A). This observation suggests that ATXN3 84Q embryos are more sensitive to Complex I inhibition, potentially reflecting heightened vulnerability in early steps of the electron transport chain (ETC).

Together, these findings highlight differential susceptibilities of ATXN3 mutant zebrafish models to ETC disruptions. While inhibition of complex IV by sodium azide affects both genotypes equally, the heightened sensitivity of ATXN3 84Q embryos to rotenone suggests specific vulnerabilities linked to Complex I inhibition and underscores the critical role of this ETC component in maintaining mitochondrial function in disease.

### Impaired Complex II function and succinate accumulation in ATXN3 84Q zebrafish and MJD mice

To explore the molecular basis for the heightened sensitivity of ATXN3 84Q zebrafish embryos to rotenone, we evaluated the levels of SDHA, a subunit of complex II in the electron transport chain, using Western blot analysis (Figure 4B). At 5 days post-fertilization (dpf), ATXN3 84Q zebrafish displayed a significant reduction in SDHA protein levels compared to WT zebrafish (Figure 4C). This decrease in complex II abundance suggests an impaired capacity to funnel electrons into the ETC through succinate oxidation.

Additionally, we quantified succinate levels in zebrafish and mice models to assess the functional consequences of reduced SDHA. Succinate levels were significantly increased in both mutant zebrafish and MJD mouse brain samples compared to their respective WT controls Figure 4 D and E). In zebrafish embryos, the succinate levels were 1.601 nmol in 23Q mutants and 2.068 nmol in 84Q mutants, representing a progressive increase in succinate accumulation with longer polyQ expansions. This trend highlights the impact of the ATXN3 mutation severity on mitochondrial function. Similarly, in mouse brain samples, succinate levels were markedly higher in the MJD mutant group compared to WT controls. WT mice exhibited succinate levels of 32.27 nmol, while the MJD mutant mice showed nearly double the amount at 64.91 nmol. These results confirm a significant disruption in mitochondrial metabolism associated with the disease phenotype in both zebrafish and mouse models.

This significant increase in succinate levels relative to WT controls is consistent with compromised activity of complex II. The observed succinate accumulation highlights a disruption in mitochondrial metabolic balance, which may exacerbate mitochondrial dysfunction in mutant zebrafish.

## Discussion

### Dysregulation of mitochondrial-associated pathways in MJD

To examine alterations to protein expression profiles in MJD brain tissue, we conducted proteomic profiling of WT and MJD transgenic mice. Our analysis revealed extensive dysregulation of mitochondrial-associated pathways, with oxidative phosphorylation (OXPHOS) being particularly affected—a result consistent with previous reports identifying mitochondrial bioenergetic deficits in polyglutamine disorders such as Huntington’s disease (HD) and spinocerebellar ataxias (SCA) (25). Similarly, other studies have reported dysregulated proteins involved in energy metabolism and mitochondrial maintenance, further supporting the involvement of mitochondrial dysfunction in these diseases. For instance, in Ki91 cerebellum mice (a mutant atxn3 knock in mice), proteins related to tricarboxylic acid cycle (TCA) and oxidative phosphorylation (OXPHOS) are dysregulated in the cerebral cortex and a substantial portion of dysregulated proteins was linked to metabolism. This dysregulation was also observed to increase with age, as metabolic-related proteins in the cerebellum showed a rise from 29% in 4-10-month-old mice to 36% in 12-14-month-old mice, suggesting a progressive worsening of mitochondrial dysfunction (26).

Quantitative pathway analysis indicated that a significant proportion of the differentially expressed proteins were downregulated (%), while % were upregulated, suggesting a net suppression of metabolic activity in MJD. Among the most affected components were complexes II and V of the ETC, which exhibited marked decreases in protein subunit abundance. Notably, Complex II (succinate dehydrogenase) is a critical bridge between the tricarboxylic acid (TCA) cycle and the ETC; its dysfunction has previously been associated with neurodegeneration and elevated ROS generation (27–30). Reduced ATP synthase (Complex V) subunits further suggest impaired energy synthesis, echoing findings in MJD patient fibroblasts and HD models (14).

Importantly, we identified similar proteomic changes across male and female cohorts of MJD mice, supporting a hypothesis that mitochondrial impairment is a robust and sex-independent hallmark of MJD pathogenesis. This finding is significant because previous studies by Almeida Et al (31) have similarly found mitochondrial dysfunction to be present in male MJD mice, but female MJD mice were not examined.

Western blot analysis of primary neurons cultured from MJD and WT mice corroborated proteomic findings. Specifically, mtND5 (Complex I) and SDHA (Complex II) levels were significantly reduced in MJD neurons. The reduction in mtND5 aligns with findings in Huntington’s disease, where intrinsic defects in Complex I subunits have been reported (32), and in Parkinson’s disease models, where pharmacological inhibition of Complex I induces oxidative stress and neuronal degeneration (33). Similarly, compromised SDHA complex II activity downregulation has been implicated in impaired electron flux and increased succinate accumulation, which can drive ROS production and activate hypoxia-like signalling (34, 35).

Pyruvate dehydrogenase downregulation further suggests upstream defects in mitochondrial substrate utilization. Pyruvate dehydrogenase downregulation has been linked to impaired neuronal glucose metabolism in Alzheimer’s disease (36).

Interestingly, we observed an upregulation of VDAC1 in MJD neurons. VDAC1 is a critical regulator of mitochondrial membrane permeability and has been associated with apoptosis driven by lipid peroxidation and reactive oxygen species (ROS) accumulation, in neurodegenerative disease models (37). The upregulation of VDAC1 in MJD neurons may therefore reflect mitochondrial vulnerability to oxidative damage, linking mitochondrial dysfunction to apoptotic pathways in disease progression. Elevated VDAC1 expression has previously been reported in Alzheimer’s disease, where it correlates with increased oxidative stress (38, 39). VDAC also influences lipid peroxidation, a key feature of ferroptosis, a form of iron-dependent cell death (40). The increased expression of VDAC1 in MJD neurons may reflect a mitochondrial stress response feedback loop, where oxidative stress exacerbates mitochondrial damage, further promoting cell death pathways such as ferroptosis.

### Abnormal mitochondrial morphology in MJD

Quantitative fluorescence microscopy revealed mitochondrial fragmentation and reduced network complexity in MJD neurons at 18 DIV. Decreased mitochondrial size, branching, and form factor are consistent with previous observations in HD, where mutant huntingtin impairs mitochondrial fusion via downregulation of MFN1/2 and OPA1 (41). Similar fragmentation patterns have been reported in MJD models, where ATXN3 mutations disrupt mitochondrial dynamics and mitophagy (15).

While overall mitochondrial number (as assessed by mtDNA qPCR) remained unchanged, the observed morphological disruptions point to compromised mitochondrial quality rather than quantity. This correlates with studies in human neuroblastoma cell line indicated no significant changes in the amount of mitochondrial biomass in cells expressing the WT (26Q) and expanded (78Q) atxn3 gene (42). This distinction is important, as recent studies have shown that mitochondrial fragmentation often precedes energetic failure and neurodegeneration in polyQ disorders (43).

### Genotype-dependent ETC dysfunction in MJD

Zebrafish embryos expressing ATXN3 84Q displayed heightened sensitivity to rotenone (complex I inhibitor) but not to sodium azide (complex IV inhibitor), suggesting a specific vulnerability to early ETC disruptions. Complex I, which plays a critical role in transferring electrons from NADH to the ubiquinone pool, is an essential entry point for mitochondrial respiration and ATP synthesis. Increased sensitivity to Complex I inhibition in ATXN3 84Q embryos may point to disruptions in mitochondrial metabolic pathways, such as altered NADH turnover or reduced complex I activity. These findings align with studies in HD zebrafish and Drosophila models, where rotenone exposure induces developmental abnormalities, ROS accumulation, and early lethality due to compromised complex I activity (44, 45).

Western blot analysis confirmed reduced SDHA protein levels in ATXN3 84Q zebrafish embryos compared to ATXN3 28Q zebrafish, while metabolite profiling revealed significant succinate accumulation in both zebrafish and MJD mouse brains. These findings are highly consistent with the literature: SDH mutations and inhibition have been shown to result in succinate buildup, which can act as an oncometabolite and pro-inflammatory signal (46–48). In neurological disease models, elevated succinate has been linked to inflammation, impaired mitochondrial respiration, increased ROS generation, and stabilization of HIF-1α under normoxic conditions, exacerbating neurodegenerative phenotypes (49–55).

The increase in succinate observed in MJD mouse brains suggests a metabolic bottleneck at complex II, which likely contributes to energy deficits and oxidative stress. When combined with rotenone treatment (complex I inhibition) in the ATXN3 84Q zebrafish embryos, the primary entry point for electrons into the ETC is blocked. In the context of reduced SDHA expression and impaired complex II function, the electron flow into the ETC is further diminished, severely limiting mitochondrial respiration and energy production. The combined inhibition of complex I by rotenone and reduced complex II capacity in the MJD zebrafish likely explains the pronounced developmental delay observed in ATXN3 84Q embryos following rotenone exposure. This suggests a critical interplay between ETC components in maintaining mitochondrial function in disease models.

The findings of this study contribute to a growing body of evidence that mitochondrial dysfunction is present when polyQ expanded ATXN3 is present, in experimental models of MJD. Findings within these experimental models, particularly the easily quantified response to rotenone treatment in the MJD zebrafish, are suitable for testing of mechanisms of dysfunction and therapeutic strategies to correct the mitochondrial and metabolic dysfunction in MJD. Further studies are needed to confirm the presence of these functional changes in human cells, and the aetiology of the proposed dysfunction.

## Notes

### Competing Interest Statement

The authors have declared no competing interest.

### Funding

- MJD Foundation: Seed Money

## References

1. Costa Mdo C, Paulson HL. Toward understanding Machado-Joseph disease. Prog Neurobiol. 2012;97(2):239–57.

2. Johri A, Beal MF. Mitochondrial dysfunction in neurodegenerative diseases. J Pharmacol Exp Ther. 2012;342(3):619–30.

3. Benarroch EE. Brain iron homeostasis and neurodegenerative disease. Neurology. 2009;72(16):1436–40.

4. Cobley JN, Fiorello ML, Bailey DM. 13 reasons why the brain is susceptible to oxidative stress. Redox Biol. 2018;15:490–503.

5. Jurcau A, Jurcau CM. Mitochondria in Huntington’s disease: implications in pathogenesis and mitochondrial-targeted therapeutic strategies. Neural Regen Res. 2023;18(7):1472–7.

6. Zhao J, Wang X, Huo Z, Chen Y, Liu J, Zhao Z, et al. The Impact of Mitochondrial Dysfunction in Amyotrophic Lateral Sclerosis. Cells. 2022;11(13).

7. Tafuri F, Ronchi D, Magri F, Comi GP, Corti S. SOD1 misplacing and mitochondrial dysfunction in amyotrophic lateral sclerosis pathogenesis. Front Cell Neurosci. 2015;9:336.

8. Mehta AR, Gregory JM, Dando O, Carter RN, Burr K, Nanda J, et al. Mitochondrial bioenergetic deficits in C9orf72 amyotrophic lateral sclerosis motor neurons cause dysfunctional axonal homeostasis. Acta Neuropathol. 2021;141(2):257–79.

9. Project Min EALSSC. CHCHD10 variants in amyotrophic lateral sclerosis: Where is the evidence? Ann Neurol. 2018;84(1):110–6.

10. Betarbet R, Sherer TB, MacKenzie G, Garcia-Osuna M, Panov AV, Greenamyre JT. Chronic systemic pesticide exposure reproduces features of Parkinson’s disease. Nat Neurosci. 2000;3(12):1301–6.

11. Sherer TB, Kim JH, Betarbet R, Greenamyre JT. Subcutaneous rotenone exposure causes highly selective dopaminergic degeneration and alpha-synuclein aggregation. Exp Neurol. 2003;179(1):9–16.

12. Kim J, Moody JP, Edgerly CK, Bordiuk OL, Cormier K, Smith K, et al. Mitochondrial loss, dysfunction and altered dynamics in Huntington’s disease. Hum Mol Genet. 2010;19(20):3919–35.

13. Kodsi MH, Swerdlow NR. Mitochondrial toxin 3-nitropropionic acid produces startle reflex abnormalities and striatal damage in rats that model some features of Huntington’s disease. Neurosci Lett. 1997;231(2):103–7.

14. Harmuth T, Weber JJ, Zimmer AJ, Sowa AS, Schmidt J, Fitzgerald JC, et al. Mitochondrial Dysfunction in Spinocerebellar Ataxia Type 3 Is Linked to VDAC1 Deubiquitination. Int J Mol Sci. 2022;23(11).

15. Hsu JY, Jhang YL, Cheng PH, Chang YF, Mao SH, Yang HI, et al. The Truncated C-terminal Fragment of Mutant ATXN3 Disrupts Mitochondria Dynamics in Spinocerebellar Ataxia Type 3 Models. Front Mol Neurosci. 2017;10:196.

16. Piasecki P, Wiatr K, Ruszkowski M, Marczak L, Trottier Y, Figiel M. Impaired interactions of ataxin-3 with protein complexes reveals their specific structure and functions in SCA3 Ki150 model. Front Mol Neurosci. 2023;16:1122308.

17. Chen H, Chan DC. Mitochondrial dynamics--fusion, fission, movement, and mitophagy-in neurodegenerative diseases. Hum Mol Genet. 2009;18(R2):R169–76.

18. Franco-Iborra S, Cuadros T, Parent A, Romero-Gimenez J, Vila M, Perier C. Defective mitochondrial protein import contributes to complex I-induced mitochondrial dysfunction and neurodegeneration in Parkinson’s disease. Cell Death Dis. 2018;9(11):1122.

19. Watchon M, Yuan KC, Mackovski N, Svahn AJ, Cole NJ, Goldsbury C, et al. Calpain Inhibition Is Protective in Machado-Joseph Disease Zebrafish Due to Induction of Autophagy. J Neurosci. 2017;37(32):7782–94.

20. Gamage H, Robinson KJ, Luu L, Paulsen IT, Laird AS. Machado Joseph disease severity is linked with gut microbiota alterations in transgenic mice. Neurobiol Dis. 2023;179:106051.

21. Cheng F, De Luca A, Hogan AL, Rayner SL, Davidson JM, Watchon M, et al. Unbiased Label-Free Quantitative Proteomics of Cells Expressing Amyotrophic Lateral Sclerosis (ALS) Mutations in CCNF Reveals Activation of the Apoptosis Pathway: A Workflow to Screen Pathogenic Gene Mutations. Front Mol Neurosci. 2021;14:627740.

22. Fath T, Ke YD, Gunning P, Gotz J, Ittner LM. Primary support cultures of hippocampal and substantia nigra neurons. Nat Protoc. 2009;4(1):78–85.

23. Link V, Shevchenko A, Heisenberg CP. Proteomics of early zebrafish embryos. BMC Dev Biol. 2006;6:1.

24. Chaudhry A, Shi R, Luciani DS. A pipeline for multidimensional confocal analysis of mitochondrial morphology, function, and dynamics in pancreatic beta-cells. Am J Physiol Endocrinol Metab. 2020;318(2):E87–E101.

25. Park J, Lee SB, Lee S, Kim Y, Song S, Kim S, et al. Mitochondrial dysfunction in Drosophila PINK1 mutants is complemented by parkin. Nature. 2006;441(7097):1157–61.

26. Wiatr K, Marczak L, Perot JB, Brouillet E, Flament J, Figiel M. Broad Influence of Mutant Ataxin-3 on the Proteome of the Adult Brain, Young Neurons, and Axons Reveals Central Molecular Processes and Biomarkers in SCA3/MJD Using Knock-In Mouse Model. Front Mol Neurosci. 2021;14:658339.

27. Zheng J, Liu S, Wang D, Li L, Sarsaiya S, Zhou H, et al. Unraveling the functional consequences of a novel germline missense mutation (R38C) in the yeast model of succinate dehydrogenase subunit B: insights into neurodegenerative disorders. Front Mol Neurosci. 2023;16:1246842.

28. Van Vranken JG, Bricker DK, Dephoure N, Gygi SP, Cox JE, Thummel CS, et al. SDHAF4 promotes mitochondrial succinate dehydrogenase activity and prevents neurodegeneration. Cell Metab. 2014;20(2):241–52.

29. Jodeiri Farshbaf M, Kiani-Esfahani A. Succinate dehydrogenase: Prospect for neurodegenerative diseases. Mitochondrion. 2018;42:77–83.

30. Laco MN, Oliveira CR, Paulson HL, Rego AC. Compromised mitochondrial complex II in models of Machado-Joseph disease. Biochim Biophys Acta. 2012;1822(2):139–49.

31. Almeida F, Ferreira IL, Naia L, Marinho D, Vilaca-Ferreira AC, Costa MD, et al. Mitochondrial Dysfunction and Decreased Cytochrome c in Cell and Animal Models of Machado-Joseph Disease. Cells. 2023;12(19).

32. Parker WD, Jr., Boyson SJ, Luder AS, Parks JK. Evidence for a defect in NADH: ubiquinone oxidoreductase (complex I) in Huntington’s disease. Neurology. 1990;40(8):1231–4.

33. Dias V, Junn E, Mouradian MM. The role of oxidative stress in Parkinson’s disease. J Parkinsons Dis. 2013;3(4):461–91.

34. Siebels I, Drose S. Q-site inhibitor induced ROS production of mitochondrial complex II is attenuated by TCA cycle dicarboxylates. Biochim Biophys Acta. 2013;1827(10):1156–64.

35. Kluckova K, Sticha M, Cerny J, Mracek T, Dong L, Drahota Z, et al. Ubiquinone-binding site mutagenesis reveals the role of mitochondrial complex II in cell death initiation. Cell Death Dis. 2015;6(5):e1749.

36. Bubber P, Haroutunian V, Fisch G, Blass JP, Gibson GE. Mitochondrial abnormalities in Alzheimer brain: mechanistic implications. Ann Neurol. 2005;57(5):695–703.

37. Shoshan-Barmatz V, Nahon-Crystal E, Shteinfer-Kuzmine A, Gupta R. VDAC1, mitochondrial dysfunction, and Alzheimer’s disease. Pharmacol Res. 2018;131:87–101.

38. Reddy PH. Is the mitochondrial outermembrane protein VDAC1 therapeutic target for Alzheimer’s disease? Biochim Biophys Acta. 2013;1832(1):67–75.

39. Yang Y, Jia X, Yang X, Wang J, Fang Y, Ying X, et al. Targeting VDAC: A potential therapeutic approach for mitochondrial dysfunction in Alzheimer’s disease. Brain Res. 2024;1835:148920.

40. Zhao Y, Li Y, Zhang R, Wang F, Wang T, Jiao Y. The Role of Erastin in Ferroptosis and Its Prospects in Cancer Therapy. Onco Targets Ther. 2020;13:5429–41.

41. Shirendeb U, Reddy AP, Manczak M, Calkins MJ, Mao P, Tagle DA, et al. Abnormal mitochondrial dynamics, mitochondrial loss and mutant huntingtin oligomers in Huntington’s disease: implications for selective neuronal damage. Hum Mol Genet. 2011;20(7):1438–55.

42. Chang JC, Wu SL, Hoel F, Cheng YS, Liu KH, Hsieh M, et al. Far-infrared radiation protects viability in a cell model of Spinocerebellar Ataxia by preventing polyQ protein accumulation and improving mitochondrial function. Sci Rep. 2016;6:30436.

43. Swinter K, Salah D, Rathnayake R, Gunawardena S. PolyQ-Expansion Causes Mitochondria Fragmentation Independent of Huntingtin and Is Distinct from Traumatic Brain Injury (TBI)/Mechanical Stress-Mediated Fragmentation Which Results from Cell Death. Cells. 2023;12(19).

44. Wang Y, Liu W, Yang J, Wang F, Sima Y, Zhong ZM, et al. Parkinson’s disease-like motor and non-motor symptoms in rotenone-treated zebrafish. Neurotoxicology. 2017;58:103–9.

45. Coulom H, Birman S. Chronic exposure to rotenone models sporadic Parkinson’s disease in Drosophila melanogaster. J Neurosci. 2004;24(48):10993–8.

46. Armstrong N, Storey CM, Noll SE, Margulis K, Soe MH, Xu H, et al. SDHB knockout and succinate accumulation are insuTicient for tumorigenesis but dual SDHB/NF1 loss yields SDHx-like pheochromocytomas. Cell Rep. 2022;38(9):110453.

47. Eijkelenkamp K, Osinga TE, Links TP, van der Horst-Schrivers ANA. Clinical implications of the oncometabolite succinate in SDHx-mutation carriers. Clin Genet. 2020;97(1):39–53.

48. Bardella C, Pollard PJ, Tomlinson I. SDH mutations in cancer. Biochim Biophys Acta. 2011;1807(11):1432–43.

49. Zhang Y, Zhang M, Zhu W, Yu J, Wang Q, Zhang J, et al. Succinate accumulation induces mitochondrial reactive oxygen species generation and promotes status epilepticus in the kainic acid rat model. Redox Biol. 2020;28:101365.

50. Huang H, Li G, He Y, Chen J, Yan J, Zhang Q, et al. Cellular succinate metabolism and signaling in inflammation: implications for therapeutic intervention. Front Immunol. 2024;15:1404441.

51. Tretter L, Patocs A, Chinopoulos C. Succinate, an intermediate in metabolism, signal transduction, ROS, hypoxia, and tumorigenesis. Biochim Biophys Acta. 2016;1857(8):1086–101.

52. Gonzalez-Meler MA, Ribas-Carbo M, Siedow JN, Drake BG. Direct Inhibition of Plant Mitochondrial Respiration by Elevated CO2. Plant Physiol. 1996;112(3):1349–55.

53. Scialo F, Fernandez-Ayala DJ, Sanz A. Role of Mitochondrial Reverse Electron Transport in ROS Signaling: Potential Roles in Health and Disease. Front Physiol. 2017;8:428.

54. Scialo F, Sriram A, Fernandez-Ayala D, Gubina N, Lohmus M, Nelson G, et al. Mitochondrial ROS Produced via Reverse Electron Transport Extend Animal Lifespan. Cell Metab. 2016;23(4):725–34.

55. Li M, Li G, Yu B, Luo Y, Li Q. Activation of Hypoxia-Inducible Factor-1alpha Via Succinate Dehydrogenase Pathway During Acute Lung Injury Induced by Trauma/Hemorrhagic Shock. Shock. 2020;53(2):208–16.

